# Comparison of cellular Na^+^ assays by flow fluorometry with ANG-2 and flame emission: Pitfalls in using ionophores for calibrating fluorescent probes

**DOI:** 10.1101/2020.01.13.904003

**Authors:** Valentina E. Yurinskaya, Nikolay D. Aksenov, Alexey V. Moshkov, Tatyana S. Goryachaya, Alexey A. Vereninov

## Abstract

Monovalent ions, sodium in particular, are involved in fundamental cell functions, such as water balance and electric processes, intra- and intercellular signaling, cell movement, pH regulation and metabolite transport into and out of cells. Fluorescent probes are indispensable tools for monitoring intracellular sodium levels in single living cells in heterogeneous cell populations and tissues. Since the fluorescence of sodium-sensitive dyes in cells is significantly different from that in an aqueous solution, the fluorescence signal is calibrated *in situ* by changing the concentration of extracellular sodium in the presence of ionophores, making the membrane permeable to sodium and equilibrating its intra- and extracellular concentrations. The reliability of this calibration method has not been well studied. Here, we compare the determinations of the intracellular sodium concentration by flame emission photometry and flow cytometry using the Na^+^-sensitive probe Asante Natrium Green-2 (ANG). The intracellular Na^+^ concentration was altered using known ionophores or, alternatively, by blocking the sodium pump with ouabain or by causing cell apoptosis with staurosporine. The use of U937 cells cultured in suspension allowed the fluorometry of single cells by flow cytometry and flame emission analysis of samples checked for uniform cell populations. It is revealed that the ANG fluorescence of cells treated with ionophores is approximately two times lower than that in cells with the same Na^+^ concentration but not treated with ionophores. Although the mechanism is still unknown, this effect should be taken into account when a quantitative assessment of the concentration of intracellular sodium is required. Sodium sensitive fluorescent dyes are widely used at present, and the problem is practically significant.

## Introduction

Monovalent ions are involved in fundamental cell functions, such as water balance and electric processes, intra- and intercellular signaling, cell movement, pH regulation and metabolite transport into and out of cells. Optical methods are indispensable tools for monitoring intracellular monovalent ion concentrations in a single living cell located in a heterogeneous cell population or tissue. Calibration of optical signals is an important problem when the intracellular ion concentration is measured by optical methods since the fluorescence intensity and spectra of ion-sensitive dyes in cells are significantly different from those in an aqueous solution. For this reason, *in situ* calibration is always performed [1-3] and is based on the variation in the extracellular concentration of the ions under consideration and the treatment of cells with ionophores, making the cell membrane permeable to these ions and likely equilibrating their intra- and extracellular concentrations [4].

It is not easy to verify the reliability of data on intracellular ion concentrations obtained in a single cell using ion-sensitive dyes by some methods because most of them require a large number of cells, and it is always questionable how uniform the cell population is in the sample. X-ray elemental analysis allows the determination of ion content in single cells with high spatial resolution, but it is very laborious and not applicable to monitoring changes in intracellular ion content in living cells. We aimed to compare data on cellular Na^+^ in human U937 lymphoma cells obtained by a known flame emission assay and by flow cytometry using the Na^+^-sensitive probe Asante Natrium Green-2 (ANG), which is currently the most sensitive and popular Na^+^-sensitive dye [5]. The effect of commonly used ionophores, gramicidin (Gram) and amphotericin B (AmB), on ANG fluorescence garnered our interested in the first place. To reveal this effect, we compared the fluorescence of ANG in cells with the same Na^+^ content in the presence and absence of ionophores. The intracellular Na^+^, tested by flame emission analysis, was changed in one case by the ionophore and in the other case by blocking the sodium pump or by inducing cell apoptosis with staurosporine (STS).

Human suspension lymphoid cells U937 were used, as they are suitable for flow fluorometry and because changes in all major monovalent ions, Na^+^, K^+^ and Cl^−^, while blocking the sodium pump and STS-induced apoptosis have been studied in detail [6-12]. We report here that the same intracellular Na^+^ concentration in ionophore-treated and untreated cells produces different ANG fluorescence signals, indicating that Gram and AmB reduce ANG fluorescence in cells. We conclude that using ANG fluorescence calibrated with ionophores does not display realistic intracellular Na^+^ when measured in the studied cells in the absence of ionophores. The decrease in ANG fluorescence in cells due to ionophores, in particular Gram, does not exclude using ANG for monitoring the relative changes in the intracellular Na^+^ concentration but leads to an error in the determination of the Na^+^ concentration in absolute units.

## Materials and methods

### Cell culture and treatment

The human histiocytic lymphoma cell line U937 was obtained from the Russian Cell Culture Collection (Institute of Cytology, Russian Academy of Sciences, cat. number 220B1). Cells were cultured in RPMI 1640 medium (Biolot, Russia) supplemented with 10% fetal bovine serum (HyClone Standard, USA) at 37 °C and 5 % CO_2_. Cells were grown to a density of 1 × 10^6^ cells per mL and treated with 10 µM ouabain (Sigma-Aldrich) or with 1 µM STS (Sigma-Aldrich) for the indicated time. Gramicidin (from Bacillus aneurinolyticus) and amphotericin B (AmB) were from Sigma-Aldrich and were prepared as stock solutions in DMSO at 1 mM and 3 mM, respectively. Na^+^-specific fluorescent dye Asante Natrium Green-2 AM ester (ANG-AM) was obtained from Teflabs (Austin, TX, Cat. No. 3512), distributed by Abcam under the name ION NaTRIUM Green™-2 AM (ab142802). Here, 5 µM Gram was added for 30 min simultaneously with ANG, and 15 µM AmB was added during the last 15 min of a 30-min ANG loading.

### Flow cytometry and cellular Na^+^ content determination

A stock solution of 2.5 mM ANG-AM was prepared in DMSO +10% Pluronic (50 µg ANG-AM was dissolved in 10 µL DMSO and mixed 1:1 with 20% (w/v) Pluronic F-127 stock solution in DMSO). After intermediate dilution in PBS, ANG-AM was added directly into the media with cultured cells to a final concentration of 0.5 µM. Incubation of cells with ANG-AM was carried out for 30 min at room temperature (∼23°C). Stained cells were analyzed on a CytoFLEX Flow Cytometer (Beckman Coulter, Inc., CA, USA). ANG fluorescence was excited using a 488 nm laser, and emission was detected in the PE channel with a 585/42 nm bandpass filter (marked below in the figures as ANG, PE). All fluorescence histograms were obtained at the same cytometer settings for a minimum of 10,000 cells. Cell population P1 was gated by FSC/SSC (forward scatter/side scatter) with a threshold set at 1×10^4^ to exclude most microparticles and debris [13] (Fig. 1A). The area (A) of the flow cytometer parameters (FSC-A, SSC-A, PE-A) was used.

**Figure 1.**
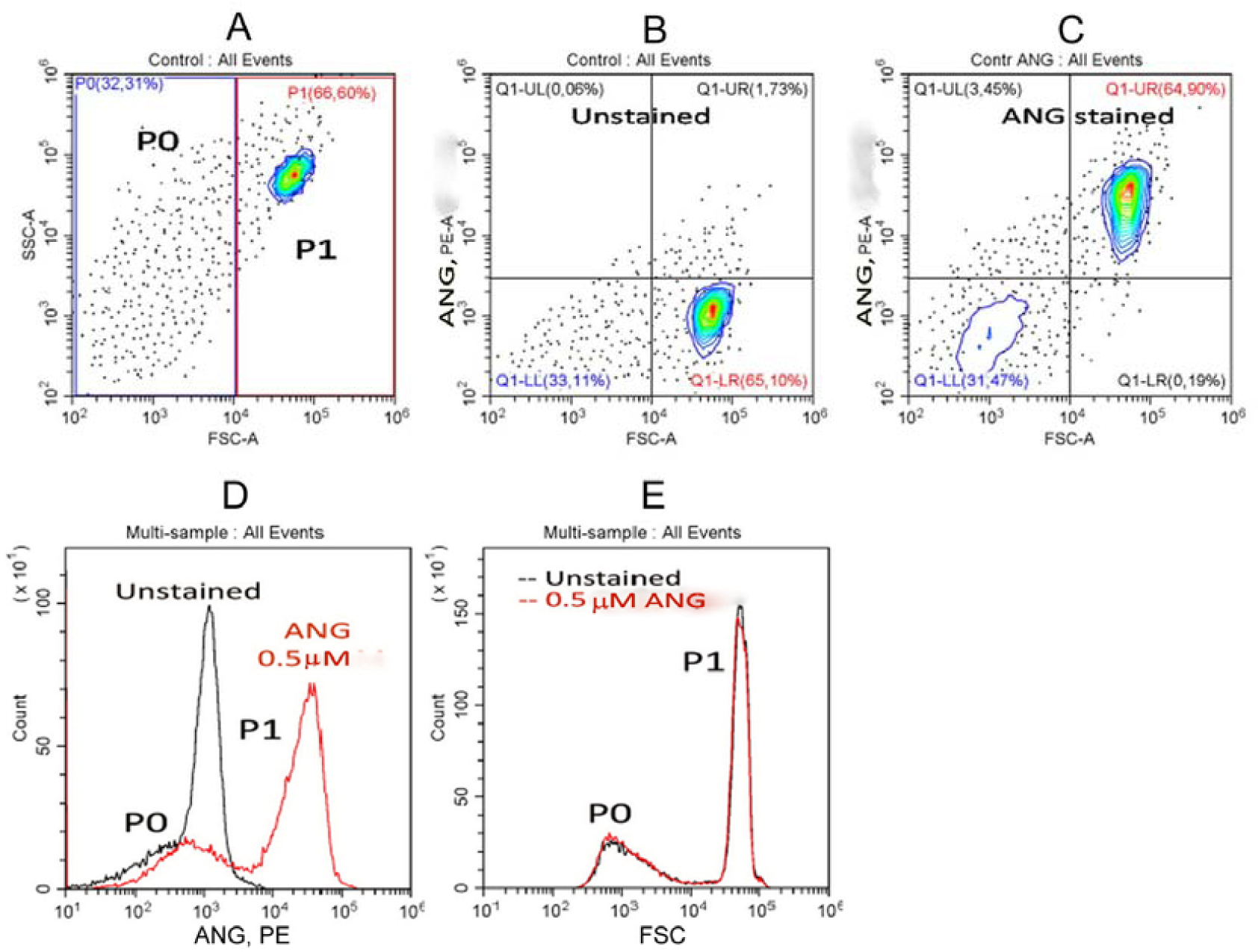
ANG fluorescence (PE), (SSC) and (FSC) distributions of normal U937 cells in unstained (A, B) and ANG-stained (C) total cell cultures. Staining was achieved by exposing cells to 0.5 µM ANG-AM for 30 min, as described in the Methods section. (D) Histograms of the fluorescence for unstained and ANG-stained cells in ANG-specific PE channel. (E) FSC histograms obtained for the same samples. The displayed data were obtained on the same day and represent at least eight separate experiments.

Cellular Na^+^ and K^+^ contents were determined by flame emission on a Perkin-Elmer AA 306 spectrophotometer as described in detail [6, 8, 10, 14, 15] and were evaluated in mmol per g of cell protein; the Na^+^ and K^+^ concentrations are given in mM per cell water. Cell water content was determined by measuring the cell buoyant density in a Percoll gradient as described earlier [13]. Cell volume was monitored additionally by a Scepter cell counter equipped with a 40-μm sensor and software version 2.1, 2011 (Merck Millipore, Germany). Cell volume, as well as cell size, changed insignificantly under all tested conditions except for the AmB treatment of cells that averaged 5.8 mL per gram of cell protein. AmB caused cell swelling.

### Microscopy

Images of ANG-stained cells were acquired on a Leica TCS SP5 confocal microscope. Cells were placed on a cover-slip in a Plexiglas holder and excited at 488 nm through a 40x Plano oil-immersion objective; fluorescence emission was collected at 500 – 654 nm. The laser power, rather than the PMT voltage, was varied to avoid image saturation for the observation of control and treated cells.

Fluorescence of hydrolyzed ANG in water solution was measured using a Terasaki multiwell plate, with a 30x water-immersion objective and a microscope equipped with an epi-fluorescent attachment. Fluorescence was excited by LED (5 W, 460 nm, TDS Lighting Co.) and passed through a dichroic filter (480-700 nm). A charged form of ANG was obtained by hydrolysis according to the recommended protocol [4]. The effect of Gram, Na^+^, and K^+^ on hydrolyzed ANG fluorescence was tested by adding 1.5 µL of a water solution of Gram (final concentration of 8 µM) with DMSO (final concentration of 0.08%), DMSO (0.08%), or H_2_O to 16.5 µL of solution in a plate well containing ANG 55 µM, KCl 125 mM, and NaCl 40 mM at pH 7.0.

### Statistical and data analyses

CytExpert 2.0 Beckman Coulter software was used for data analysis. The means of PE-A for appropriate cell subpopulations given by CytExpert software were averaged for all samples analyzed. Data were expressed as the mean ± SD for the indicated numbers of experiments and were analyzed using Student’s t test. P < 0.05 was considered statistically significant.

## Results

### P1 and P0 subsets in ANG and FSC (Forward Scatter) flow diagrams of the total U937 population

Flow cytometric analysis of the original U937 cell culture shows two main particle subsets in FSC/SSC (Forward Scatter/Side Scatter) and FSC/ANG, PE (PhycoErythrin) plots that we denote as P0 and P1 (Fig. 1A-C). P1 represents intact cells, while P0 contains particles, vesicles or debris [13]. The cell culture as a whole is analyzed using the flame emission method, and it is usually unknown which part consists of cells and which consists of the fragments of cells and debris. Flow cytofluorometry with ANG provides the answer to this question. Fluorescence intensity plots contain the P0 and P1 subsets similar to those in FSC histograms but with slightly broader peaks (Fig. 1D, E). A comparison of Fig. 1B and 1C shows that ANG fluorescence (PE-A signal) increases noticeably for all P1 cells and only slightly for the P0 subset. Flow cytometry of ANG-stained U937 cells allows evaluation of the uniformity of cell populations used in the flame emission assays. The cumulative ANG signal associated with P1 (calculated from the mean fluorescence and the number of particles) accounts for approximately 98% of all the ANG contained in P1+ P0. The typical P0/P1 ratio for the integral ANG signal was 0.019±0.001 (n = 8). A similar ratio for the FSC signal was 0.023±0.002. This means that the sodium content reported by the flame emission analysis corresponds to the P1 subset. Therefore, we further consider only ANG fluorescence of the P1 subset. Microscopy confirms the uniform distribution of ANG staining in the P1 cell population and a certain heterogeneity in the distribution of ANG within the cells (Fig. 2). Flow cytometry has an advantage over fluorescence photometry using microscopy due to the higher accuracy of single cell photometry and the high statistics (10-20 thousand cells).

**Figure 2.**
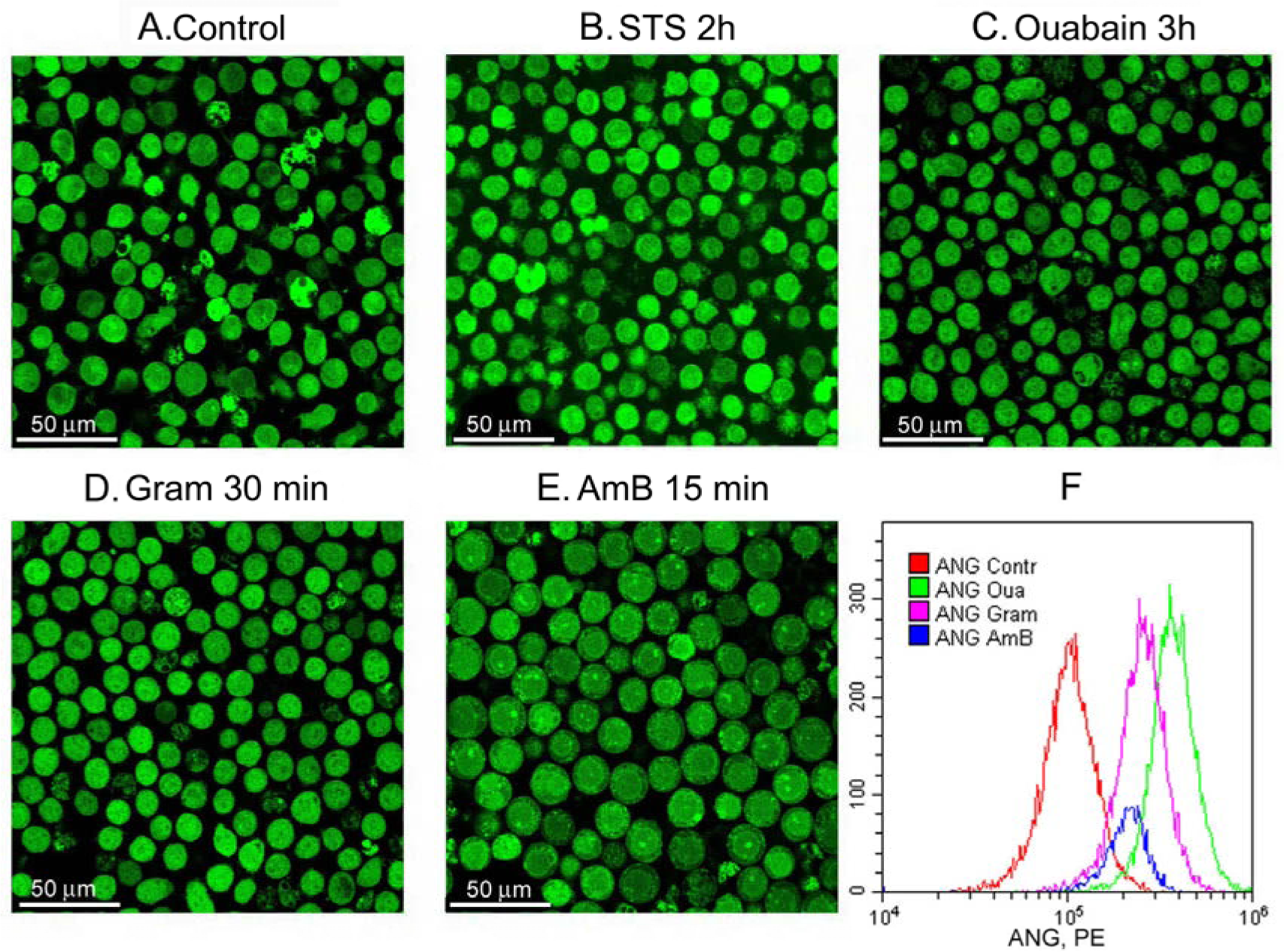
Microscopy of ANG-stained cells treated with STS, ouabain, Gram or AmB. U937 cells were treated with STS (B) or ouabain (C) during the indicated time and were stained with ANG-AM for the last 30 min, as described in the Methods section. Alternatively, cells without treatment with ouabain or STS were stained with ANG-AM in the presence of Gram (D) or AmB (E) during the last 15 min of the 30-min ANG-AM staining process. After staining, cells were analyzed on a flow cytometer except cells treated with STS (F) or viewed on a confocal microscope (A-E). Fluorescence images were acquired on the microscope at a laser power of 100% for the control and STS-treated cells, 30% for ouabain-treated cells, 40% for gramicidin-treated cells and 70% for AmB-treated cells. Flow cytometer histograms (F) were obtained at the same cytometer setting and the same loading dye concentration and can be used for comparison of the fluorescence intensities in cells under different conditions in A, C-E.

Cell loading was tested at several concentrations of ANG (Fig. 3A). The concentration of 0.5 µM was chosen as sufficient because ANG fluorescence of the P1 subset significantly overlaps the background cell autofluorescence (Fig. 1D, 3D). The protocol recommended by the manufacturer calls for loading of ANG for 30 min followed by a wash step and additional incubation to allow complete deesterification of the dye. This procedure has often been used to prepare cells for microscopic observation, but we found that washing of cells to remove the external ANG had no significant impact on the ANG signal measured on a flow cytometer (Fig. 3C). Although the presence of serum in the loading medium decreased the cell fluorescence, the signal remained sufficiently strong (Fig. 3C, 3D), and we preferred to keep the serum. Thus, a 30-min incubation with 0.5 µM ANG in normal media with serum without a subsequent wash step was used in all experiments. Importantly, this staining procedure did not affect the FSC/SSC histogram, which is known to be a sensitive indicator of cell health (Fig. 1E).

**Figure 3.**
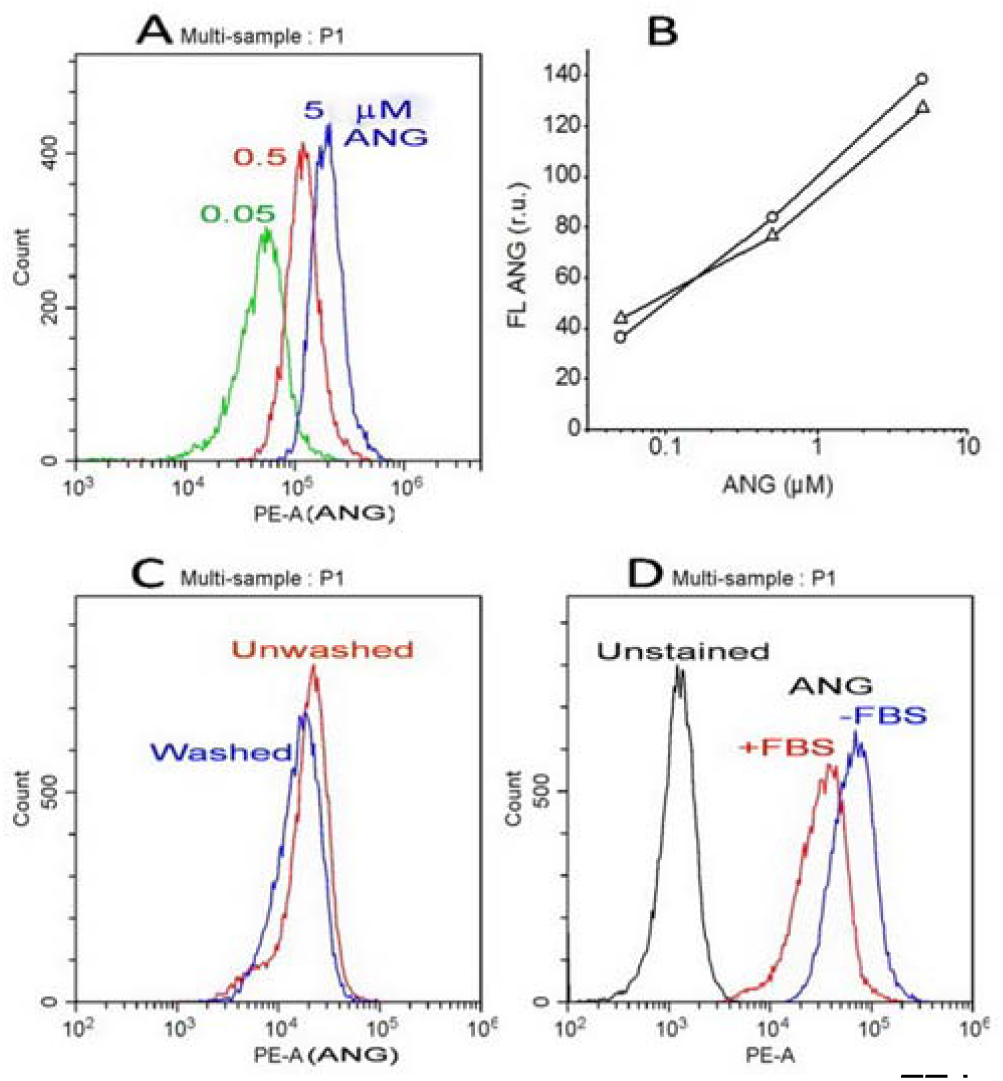
Effects of ANG concentration (A, B), washing (C) and serum (D, FBS) on ANG fluorescence in the P1 cell subset. The displayed histograms were obtained in independent experiments with identical instrument settings. (B) Dependence of cell ANG fluorescence on ANG-AM concentration in serum-free RPMI media, for 2 experiments, as shown in Figure 3A.

### Comparison between [Na^+^]_in_ estimation by ANG fluorescence and by flame emission assay

Ionophores are commonly used to calibrate fluorescent ion probes by controlling the intracellular ion concentration. In this study, we did not need the usual step of calibration, and we applied a different approach. We compared the relative changes in cellular sodium, as measured by flame emission analysis, with the relative changes in ANG-Na fluorescence detected by a flow cytometer, which reflects the total fluorescence of a single cell, averaged over 20 thousand measurements. An increase in cellular Na^+^ was induced by stopping the pump with ouabain or by STS; the effects of these stimuli on cell ion composition have been previously characterized in detail [6-13]. The treatment of cells with Gram commonly used in ANG calibration gave another way to increase cellular Na^+^. Indeed, both ouabain and Gram increased ANG fluorescence significantly (Fig. 4A and B). The cell volume measured by a Scepter cell counter practically did not change in cells treated with ouabain or gramicidin for the indicated time (Fig. 5). The FSC histograms do not change in these cases as well (Fig. 4C and D). FSC is usually considered an indicator of cell size, albeit with some reservations [13, 16, 17].

**Figure 4.**
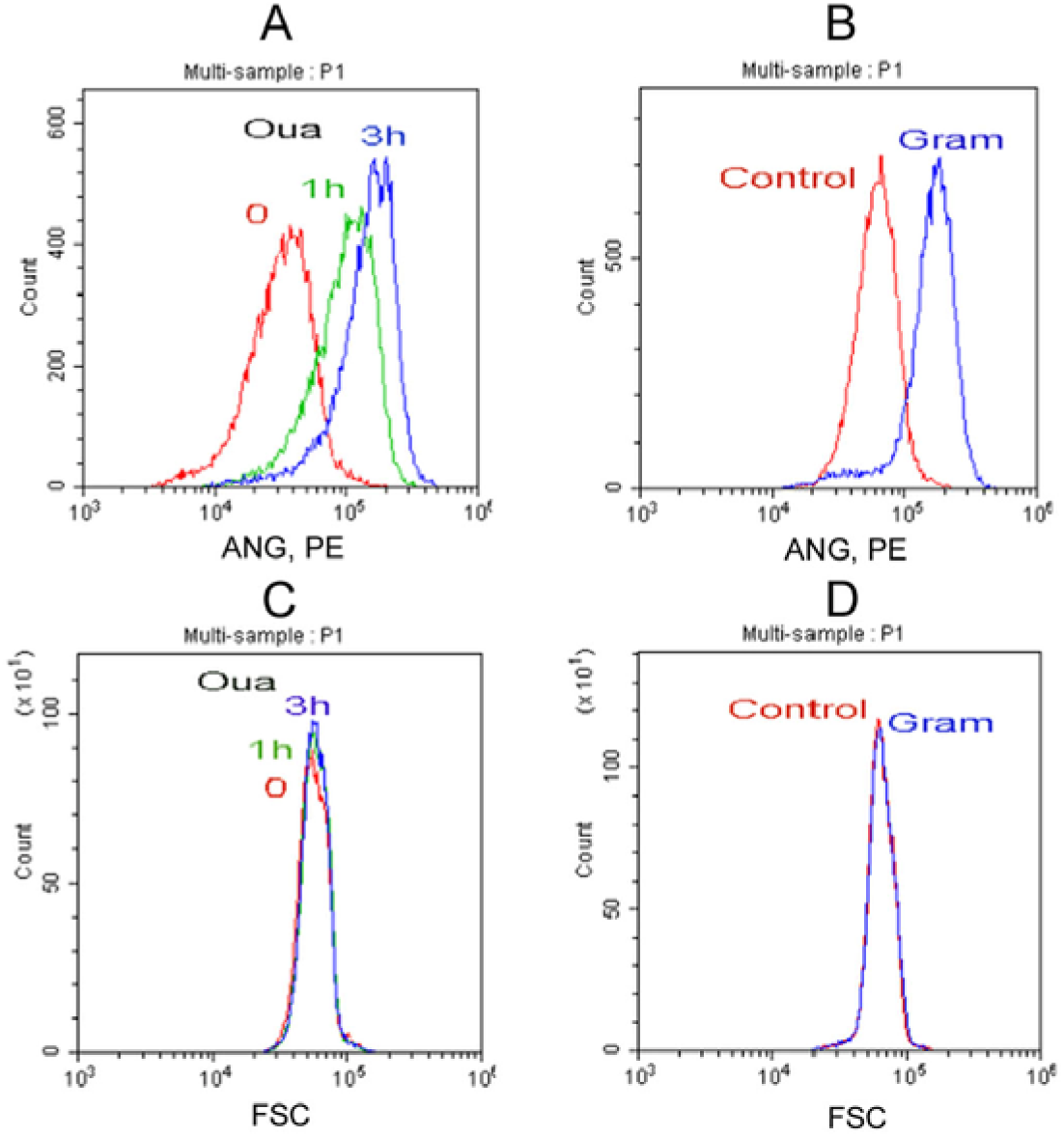
ANG fluorescence (A, B) and FSC-histograms (C, D) for the ouabain- and gramicidin-treated cells. Histograms were obtained in independent representative experiments with the same instrument settings for the compared data. Cells were treated with 10 µM ouabain for the indicated time; 5 µM gramicidin was added for 30 min with ANG-AM.

**Figure 5.**
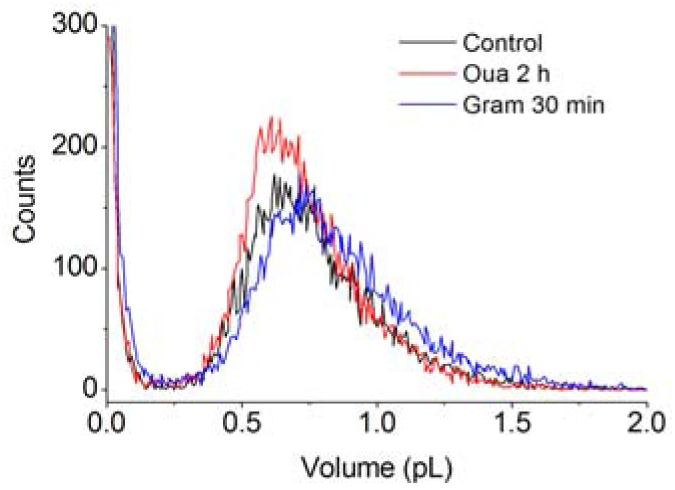
Cell volume histograms obtained by a Scepter cell counter for cell cultures treated with 10 µM ouabain or 5 µM Gram for the indicated times. Histograms for comparison are obtained in the same representative experiment.

Flow cytometry showed an excellent agreement in the changes in fluorescence and in the cellular Na^+^ increase due to stopping the pump with ouabain and STS-induced apoptosis (Fig. 6A and B). Gram and AmB added to U937 cells increased the intracellular Na^+^ to a greater extent than did ouabain for 3h, as obtained by flame emission assay. However, the increase in ANG fluorescence was much smaller for AmB and Gram than for ouabain (Fig. 6A). The discrepancy between the fluorescence of ANG and the Na^+^ concentration by flame emission assay in the presence of ionophores is especially striking when comparing the data on the same graph (Fig. 6C). It should be noted that all compared flow cytometry data were obtained with the same instrumental settings and were well reproduced. We conclude that ANG fluorescence does not display a realistic cellular Na^+^ concentration if the fluorescence is measured in the absence of ionophores while it was calibrated in its presence.

**Figure 6.**
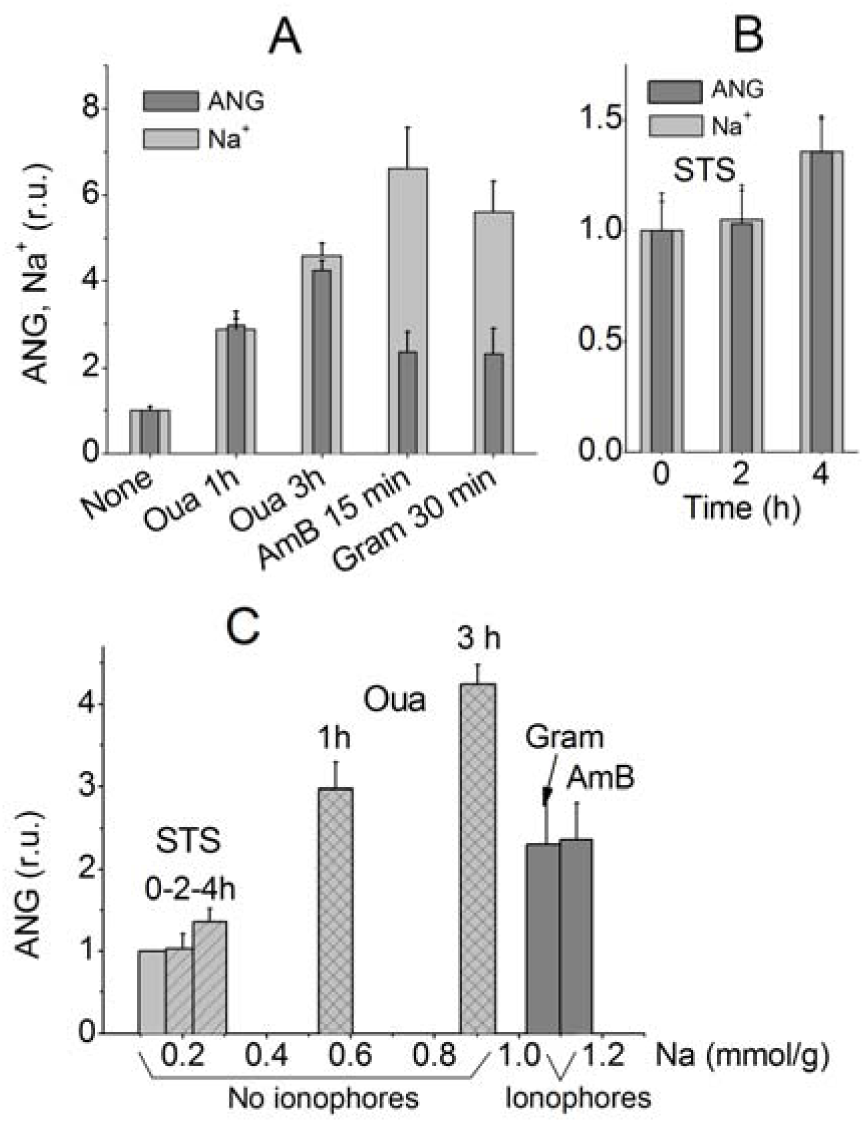
Comparison of changes in ANG fluorescence and cellular Na^+^ by flame emission assay under various conditions. (A) Cells were treated with 10 µM ouabain, 5 µM Gram, or 15 µM AmB for the indicated times; (B) cells were treated with 1 µM STS to induce apoptosis; (C) ANG fluorescence at different Na^+^ concentrations obtained with and without ionophores. Other indications are on the graphs. Means ± SD were normalized to the values in untreated cells for 3-5 experiments (in duplicate for cellular Na^+^ content). The measured Na^+^ content in mmol/g cell protein corresponds to intracellular concentrations ranging from 34 to 154 mM.

### Gramicidin does not influence ANG fluorescence *in vitro*

We tested the effect of Gram on hydrolyzed ANG. ANG fluorescence was measured using a Terasaki multiwell plate and a fluorescent microscope, as described in the Methods section. The ANG-AM used for probing intracellular Na^+^ is practically nonfluorescent *in vitro* (Fig. 7). It is believed that after penetrating the cell membrane, ANG-AM is hydrolyzed by non-specific esterases. Indeed, after hydrolysis of ANG-AM *in vitro* by a known protocol [4], ANG strongly responds to Na^+^ but responds relatively weakly to K^+^, supporting the view that the same can occur in cells (Fig. 7). The most important finding with regard to our problem was the lack of any quenching effect of Gram on the fluorescence of hydrolyzed ANG at ion concentrations imitating those in the cytoplasm.

**Figure 7.**
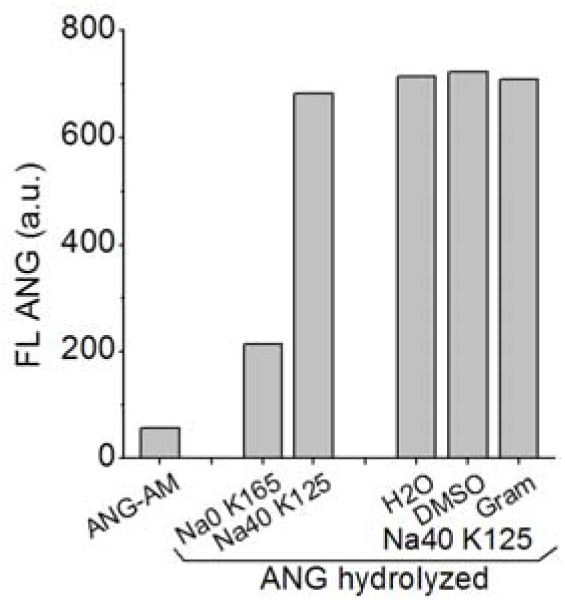
The effect of Na^+^, K^+^ and Gram on the fluorescence of ANG-AM and on hydrolyzed ANG in water solutions. The values are the averages for 2 measurements. The data from one representative experiment of two are shown.

## Discussion

ANG-2 is currently the most commonly used and sensitive probe to study sodium distribution and dynamics in various types of cells. ANG has been used in brain slices, neurons and their dendrites and axons, astrocytes [18-24], HEK293 cells [23], cockroach salivary gland cells [26], cardiomyocytes [27], and prostate cancer cell lines [5]. Optical Na^+^ monitoring is indispensable in neuroscience because of the extreme complexity of the object structure. The number of studies using fluorometric Na^+^ measurements is rapidly increasing [28]. To our knowledge, our study is the first to use ANG in flow cytometry and the first to compare Na^+^ data obtained using flow fluorometry and flame emission assays. Using flow fluorometry with ANG enables testing of cell samples used for flame emission assays to verify that they consist of relatively homogeneous cell populations. Thus, the data obtained by the optical method on a single cell can be compared with the average data found by flame emission on the cell population.

It is known that the complex of ANG with Na^+^ in cells differs from the analogous complex in a water solution according to the K_d_ and other properties [5, 18]. Similar differences were found earlier for other ion-specific probes, e.g., for SBFI [1] and Sodium Green [2]. It was reported that K^+^ can change the dependence of the sodium probe fluorescence on the Na^+^ concentration [5, 29]. Harootunian and colleagues in their pioneering work [1] supposed that SBFI spectral characteristics in the cytoplasm could be different in cells and in external water media due to differences in viscosity. It should be noted that Rose and colleagues [30] did not observe a significant influence of the differences in viscosity on CoroNaGreen fluorescence.

Because of differences in the properties of probes inside cells and in water solutions, *in situ* calibration is always performed, which is based on the variation in the extracellular concentration of the ions under consideration and the treatment of cells with ionophores, making the cell membrane permeable to Na^+^ and likely equilibrating its intra- and extracellular concentrations. Our study is one of the first to compare the data on intracellular Na^+^ contents obtained by fluorometry and those obtained by a well-established flame emission assay. Cellular Na^+^ was altered in three different ways: by stopping the sodium pump with ouabain, by inducing apoptosis with staurosporine, and by treating cells with ionophores. We found that ANG fluorescence in cells treated with gramicidin or amphotericin was approximately two-fold lower than that in cells with the same sodium concentration but without ionophores. Our tests of the *in vitro* interactions of hydrolyzed ANG with Gram lead to the conclusion that a decrease in ANG fluorescence in cells caused by Gram is unlikely due to the “simple” physical quenching of ANG in cells. It could be assumed that the effect of ionophores on ANG fluorescence in the cell is determined by changes in the interaction of ANG with some unknown cytoplasm components or by an influence of ionophores on the process of “de-esterification” of the ANG-AM in cells.

In view of the gramicidin effect on ANG fluorescence observed in our experiments, the calibration process without ionophores is interesting. Rose and colleagues compared traditional calibration methods using gramicidin with the calibration without gramicidin by directly dializing single cells via a patch pipette by saline with different Na^+^ contents [31]. They stated that similar results were obtained. It should be noted that the SBFI probe in the ratiometric mode was used in this study.

An interesting question concerns equalizing extra- and intracellular concentrations of cations in cells treated with Gram and other ionophores. Our flame emission assay showed that there was no exact equalization of monovalent cation concentrations in the medium and in cells after treatment with Gram. The Na^+^ level becomes somewhat higher than that in the medium, while the K^+^ level is approximately 25 mM at an external concentration 5 mM after a 30-min treatment of U937 cells with 5 µM gramicidin (Fig. 8).

**Figure 8.**
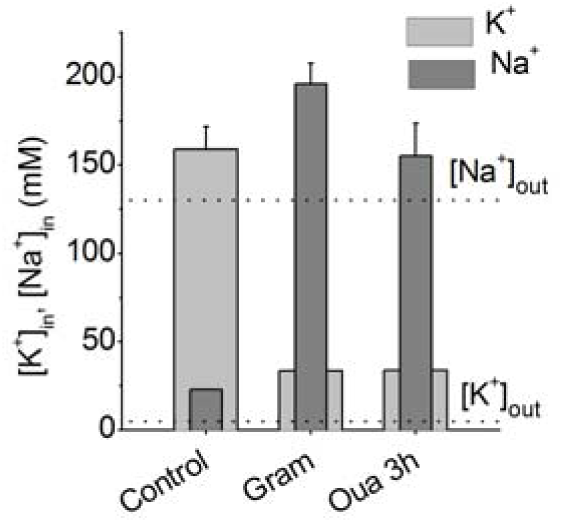
[K^+^]_in_ and [Na^+^]_in_ changes in U937 cells treated with 5 µM Gram for 30 min or with 10 µM ouabain (Oua) for 3 h. Data obtained by flame emission assay. Mean ± SD from 3 experiments with duplicate determinations. Dotted lines indicate [Na^+^]_out_, [K^+^]_out_

It is believed that in the media where Cl^−^ is partly replaced with gluconate, Donnan’s rules become insignificant [1]. We calculated changes in the K^+^ and Na^+^ concentrations using our recently developed computational tool based on Donnan’s rules [9] for model cells with parameters similar to U937 cells. An increasing cell membrane permeability for K^+^ and Na^+^ (pK and pNa in the calculation, respectively) by approximately 10 and 50 times was choosed, imitating the gramicidin effect. The K^+^ level in the cell decreased but remained higher than the extracellular level, while Na^+^ increased to a level slightly higher than the extracellular level (Fig. 9). The calculation showed that the chloride permeability of the cell membrane is the most important determinant of the rate of the monovalent ion redistribution after blocking the Na/K pump or after an increase in Na^+^ and K^+^ permeabilities due to ionophores. For this reason, a long time is required for the “standard” cell, such as U937, to swell after treatment with ouabain or Gram, and much less time is required after treatment with amphotericin, which increases not only Na^+^ and K^+^ but also Cl^−^ permeability of the cell membrane.

**Figure 9.**
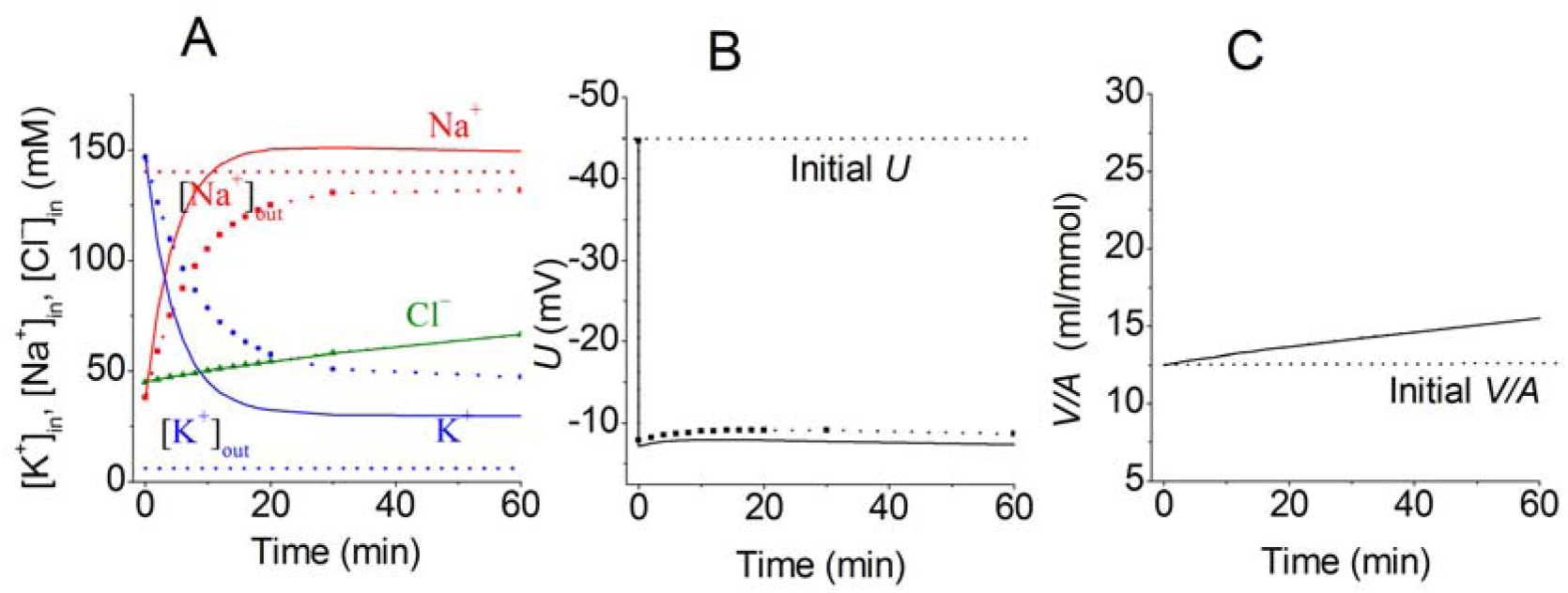
[K^+^]_in_, [Na^+^]_in_, and [Cl^−^]_in_, membrane potential (*U*) and cell volume (*V/A*) calculated for model cells with parameters similar to those of U937 cells. In the balance state (t<0), in mM [Na^+^]_in_ 38, [K^+^]_in_ 147, [Cl^−^]_in_ 45, V/A 12.50 pNa 0.00382 pK 0.022 pCl 0.0091; Gram is added at t=0 and pNa, pK are set 0.1 (dotted lines with symbols) or 0.2 (solid lines). The other parameters remain unchanged. Dashed lines indicate [Na^+^]_out_, [K^+^]_out_ (A) and the initial parameters (B, C). Cell volume is given in mL per mmol of impermeant intracellular anions (*V/A*).

The problem of equalizing intra- and extracellular Na^+^ concentrations by ionophores is discussed by Boron and colleagues, who were among the few who performed flame emission analysis in parallel with the determination of intracellular Na^+^ in HeLa cells using an SBFI probe with various ionophores and microscopy cell imaging [3]. They found that “no two of the three ionophores (gramicidin, nigericin, monensin) were adequate to totally permeabilize the cells to Na” and “the combination … does in fact equalize”. With respect to these results, we should note that emission analysis of the monolayer of HeLa cells is highly dependent on the estimation of the amount of extracellular sodium in the sample, and using inulin does not guarantee the real situation of intracellular sodium.

### Our general conclusion

Fluorescence of the sodium-sensitive dye ANG measured in the studied cells in the absence of ionophores does not display a realistic intracellular Na^+^ concentration when calibrated using ionophores but allows monitoring of the relative changes in the intracellular Na^+^ concentration.

## Acknowledgments

The authors are grateful to Dr. Michael Model (Department of Biological Sciences, Kent State University, Kent, Ohio 44242, USA) for manuscript revision and suggesting improvements. The research was supported by the State assignment of Russian Federation (No. 0124-2019-0003).

## Author contributions

All authors contributed to the design of the experiments, performed the experiments, and analysed the data. AV wrote the manuscript with input from all authors. All authors have approved the final version of the manuscript and agree to be accountable for all aspects of the work. All persons designated as authors qualify for authorship, and all those who qualify for authorship are listed.

